# OfWRKY33 binds to the key linalool synthase gene *OfTPS7* to promote linalool synthesis in *Osmanthus fragrans* flowers

**DOI:** 10.1101/2024.10.17.618933

**Authors:** Wan Xi, Meng-Yu Jiang, Lin-lin Zhu, Xu-Mei Zeng, Huan Ju, Qin-Lian Yang, Ting-Yu Zhang, Cai-Yun Wang, Ri-Ru Zheng

## Abstract

Volatile aroma compounds make significant contributions to human perception of flowers. *Osmanthus fragrans* is a famous aroma plant and linalool along with its oxides are proved to be the dominant aroma active compounds. Although some terpene synthases (TPSs) have been characterized, a comprehensive study of the hub metabolic gene and its transcriptional regulation remain to be revealed. Here, we selected a specific cultivar Boyeyingui (BBYG) with the highest content of linalool among 20 wide-cultivated cultivars for genome and transcriptome sequencings. Among the 25 new putative *OfTPSs,* only *OfTPS6*, *OfTPS7* could exclusively produce linalool *in planta*. Biochemical analysis demonstrated that OfTPS6, OfTPS7 were able to catalyze geranyl diphosphate (GPP) into linalool and a small proportion of other monoterpenes *in vitro*. Spatial and temporal correlation analysis further confirmed the transcript level of *OfTPS7* was closely associated with linalool content in diverse cultivars and different tissues, suggesting *OfTPS7* was the essential linalool synthase gene. Combined with yeast one-hybrid screen and weighted correlation network analysis (WGCNA), a nucleus-localized transcriptional factor OfWRKY33 was considered as a potential modulator. Y1H, dual-luciferase assay and electrophoretic mobility shift assay (EMSA) demonstrated that OfWRKY33 directly bound to the W-box of *OfTPS7* promoter to stimulate its transcription. OfWRKY33 could coordinately induce the expressions of *OfTPS7*, *1-deoxy-d-xylulose 1* (*OfDXS1*), thereby promoting the linalool formation. The results first revealed the hub linalool synthase gene *OfTPS7* and a novel TF participating in the complex transcriptional regulation of linalool biosynthesis in *O. fragrans* flowers.

## Introduction

Aroma compounds make great contribution to consumers’ perception of flowers. Terpenoids are the dominant aroma compounds in diverse plants (Stirling et al., 2024; Bergman et al., 2024). hey not only play vital roles in development and interaction with environment for plants (Pichersky and Raguso, 2018), but also greatly contribute to horticulture, perfumery, therapeutics and flavorings (Li et al., 2019; Bao et al., 2020; Wu et al., 2022). Current studies mainly focus on world-wide fruits, vegetables and flowers (Zhou et al., 2020; Wei et al., 2021; Shang et al., 2024), researches on more characteristic species can expand understanding of terpenoid biosynthesis and regulation.

*Osmanthus fragrans* is a famous aroma plant which can be traced back for more than 2,500 years in China. Terpenoids comprise of approximately 70% of total aroma compounds in most cultivars (Zheng et al., 2017; Fu et al., 2019). Linalool and its oxides are important aroma active compounds in flowers of diverse *O. fragrans* cultivars, imparting essential floral notes to fresh flowers and its essential oil (Wang et al., 2017; Wu et al., 2022). Although 40 *OfTPSs* were annotated in *O. fragrans* (ORF) genome and 7 *OfTPSs* were highly expressed in flowers compared with other tissues, functional identification was lacking to further elucidate their roles (Yang et al., 2018). In our previous studies, *OfTPS1/2/5* have been screened by homologous sequence alignment with *A. thaliana* and functionally characterized as linalool synthase genes (Zeng et al., 2016). However, *OfTPS2* can only be detected in 2 cultivars, *OfTPS1* and *OfTPS5* have low transcript levels in flowers of some cultivars. These results imply that the key linalool synthase genes have not been fully illustrated yet. Although plants possess a considerable number of *TPSs* in the genome, a limited proportion of TPSs are capable of producing terpenoids (Chuang et al., 2018; Zhou et al., 2020). For instance, in *Freesia × hybrida*, FhTPS1 is responsible for the formation of linalool, while FhTPS4, FhTPS6, and FhTPS7 are bifunctional enzymes producing mono-/sesqui-terpenes simultaneously (Gao et al., 2018). These results suggested that genome sequencing combined with functional identification both *in vivo* and *in vitro* would provide a comprehensive view of terpenoid biosynthesis in plants (Zhou et al., 2020; Li et al., 2024).

In recent years, transcription factors (TFs) including MYB, AP2/ERF, bHLH, MADS and WRKY have been reported to regulate the terpenoid biosynthesis by directly binding to the promoters of important pathway genes (Yang et al., 2020; Gong et al., 2021; Yang et al., 2021; Wei et al., 2022; Zhao et al., 2023; Chen et al., 2024). FhMYB21L2 mediated linalool synthesis by binding to the MYBCORE site on the promoter of *FhTPS1*(Yang et al., 2020). OfMYB21 could bind to the promoter of *OfTPS2* and positively affected linalool synthesis (Lan et al., 2023). WRKY TFs, one of the most essential families of plant-specific TFs, are extensively involved in plant growth and development by recognizing the cis-acting element W-box (Zhao et al., 2023; Chen et al., 2024). Recently, WRKY TFs have also been reported to participate in the secondary metabolite synthesis and regulation. In *Salvia miltiorrhiza,* overexpression of *SmWRKY1* could significantly increase the content of tanshinone (Cao et al., 2018). *CrWRKY42* promoted carotenoid accumulation by inducing the expression of multiple carotenoid biosynthetic genes (Chen et al., 2024). 154 WRKY genes were screened with WRKY domains in *O. fragrans* genome and 8 WRKY genes presented flower-specific expression patterns (Ding et al., 2020). However, the transcriptional regulation of aroma compounds in *O. fragrans* flowers remain elusive. Here, a comprehensive analysis was performed to clarify the key linalool synthase gene along with its regulation mechanism. First, a typical cultivar ‘BBYG’ with highest content of linalool among 20 cultivars has been selected and subjected to genome analysis and transcriptome sequencing. The functions and expression patterns of all the *OfTPSs* were determined, *OfTPS7* was confirmed to play crucial role in linalool formation in most cultivars. The TF *OfWRKY33* was screened by Y1H assay using *OfTPS7* as a bait combined with weighted correlation network analysis (WGCNA) derived from transcriptome sequencings. *OfWRKY33* could stimulate transcription of *OfTPS7* by directly binding to the W-box in the promoter and finetuned multiple MEP pathway genes, thereby promoting linalool biosynthesis. Our findings first illustrate the key linalool synthase gene together with its complex transcription regulation in *O*. *fragrans* flowers.

## Results

### Linalool and its oxides are key characteristic aroma compounds of *O. fragrans* flowers

Headspace solid phase microextraction (HS-SPME) combined with GC-MS (gas chromatography-mass spectrometry) was applied to determine the volatile aroma compounds in the full blossoming flowers of 20 wide-cultivated *O. fragrans* cultivars (Fig1A). Terpenoids, phenylpropanoids, fatty acid derivatives and other aroma compounds were found. Terpenoids were the most abundant aroma compounds in all the cultivars (Fig 1B). Linalool and its oxides were the dominant aroma active compounds due to its high contents and odor activity values (OAVs, Supplementary Table 1). They could impart noticeable floral fragrance to *O. fragrans* flowers. The specific cultivar ‘BYYG’ was subjected for further study of linalool biosynthesis due to its maximum linalool content among 20 cultivars (Fig 1C). Linalool and its oxides accounted for 48% and 7.4% of the terpenoids in ‘BYYG’, respectively (Fig 1D). In addition, temporal analysis demonstrated that linalool and its oxides increased from bud stage and achieved its maximum content in the full blossoming stage. Thus, our study selected the specific cultivar ‘BYYG’ for genome and transcriptome profiles in an attempt to elucidate the complex linalool biosynthesis in *O. fragrans* flowers.

**Figure 1.**
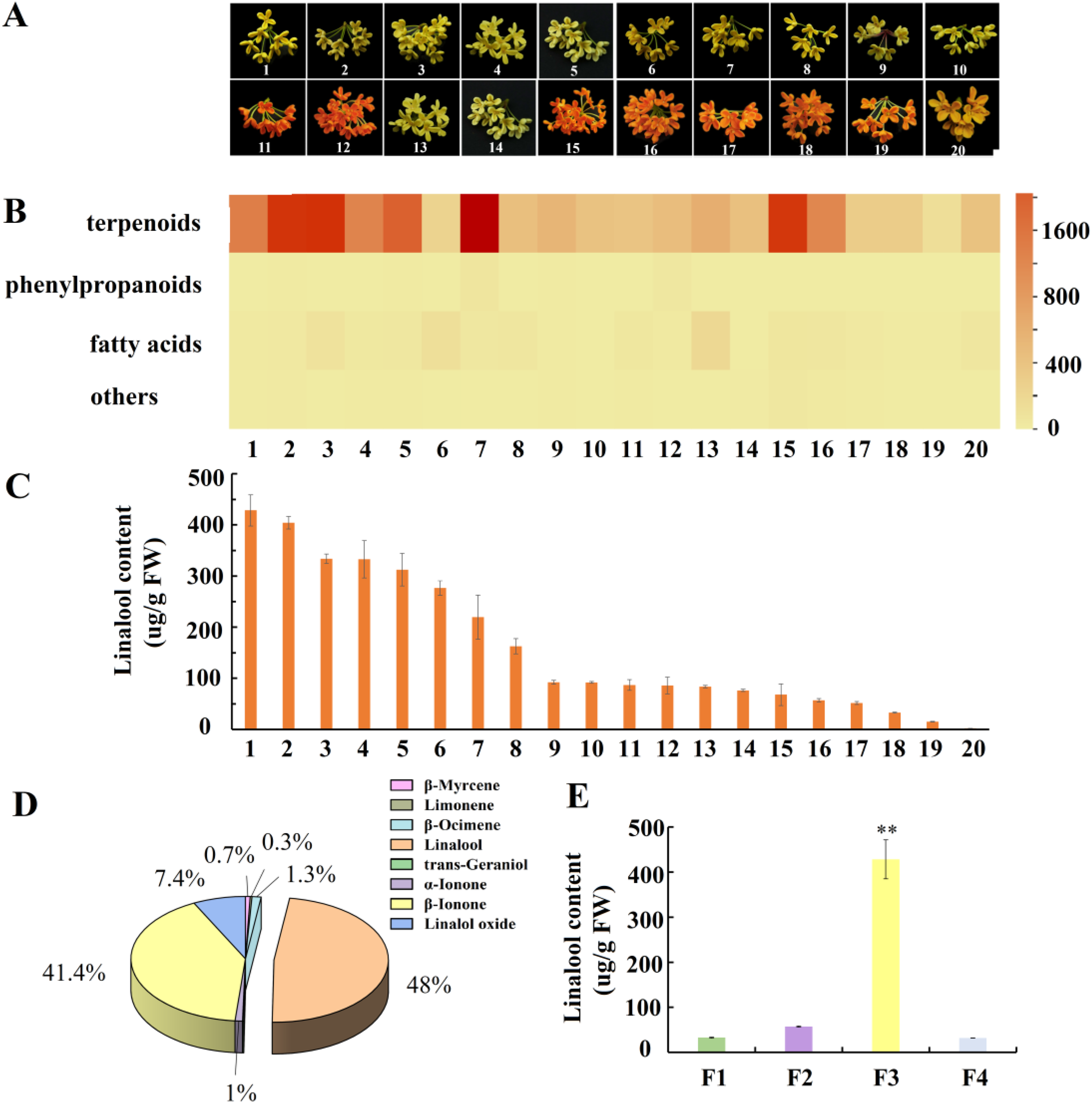
Volatile aroma compounds in *O. fragrans* flowers. A, Full blossoming flowers of 20 wide-cultivated *O. fragrans* cultivars; 1: BBYG, 2: MJYG, 3: DZ, 4: DYH, 5: FDZH, 6: SY, 7: ZY, 8: RYJG, 9: GZSJG, 10: JQZ, 11: FJH, 12: LZDG,13: HCJG, 14: LYYG, 15: JR, 16: HYNX, 17: ZSG, 18: XSDG, 19: ZYH, 20: CHDG. B, Heatmap of different volatile aroma compound contents in 20 cultivars; C, Linalool contents of the full blossoming stage flowers in 20 cultivars; D, Proportion of different terpenoids in full blossoming flowers of BBYG; E, Linalool contents of the four blossoming stages of BBYG, F1: bud stage; F2: initial blossoming stage; F3: full blossoming stage; F4: late blossoming stage. Statistical significance with respect to the reference sample (control) was determined by the Student’s *t*-test. *\*P<0.05, **P<0.01, ***P<0.001*

### Identification of the terpene synthase gene family

To initiate the analysis of the *OfTPSs* family in *O. fragrans*, ‘BYYG’ flowers with the highest content of linalool were subjected to genome sequencing and transcriptome sequencing of four blossoming stages (Fig 1B). We used a combination of sequencing technologies, including PacBio, SMRT, and Hi-C, to construct a refence genome for *O. fragrans.* A total of 264.92 Gb HiFi clean data and 49.86 Gb SMRT clean data were obtained, respectively. The genome size was estimated to be 711.42 Mb with a high heterozygosity of 1.23% according to *K*-mer-based statistics. The assembled genome was 726Mb with a contig N50 of 18.83Mb (Table 1, Supplementary Table 2), significantly longer than that in ‘Liuyeyingui’ (OFL, contig N50 =2.36 Mb) (Chen et al., 2020) and ‘Rixianggui’ (OFR, contig N50=1.60 Mb) (Yang et al., 2018). We applied Hi-C sequencing technology and anchored 98.21% of the sequences onto 23 chromosomes (Fig 2A, B). Based on this high-quality genome assembly, the genome completeness reached 98.20% evaluated by BUSCO. The gene predictions covered 97.50% of highly conserved core proteins in the eudicot lineage, which was higher than those of other two *O. fragrans* genomes (94.50% and 96.80%, respectively). High mapping rate of ISO-seq reads and the high BUSCO score suggested the high completeness and accuracy of the assembled genome. In summary, the evaluations demonstrated that the *O. fragrans* ‘BYYG’ genome (OFB) had superior quality in assembly and annotation. Based on the 1genome, transcriptome sequencing of four developmental stages of flowers were conducted as well. After library construction, Illumina sequencing and assembly, approximately 31.81, 30.8, 33.55 and 33.35 million total clean reads for four samples were generated, respectively (Supplementary Table 3), the above data indicates that the transcriptome data was qualified for subsequent analysis.

**Figure 2.**
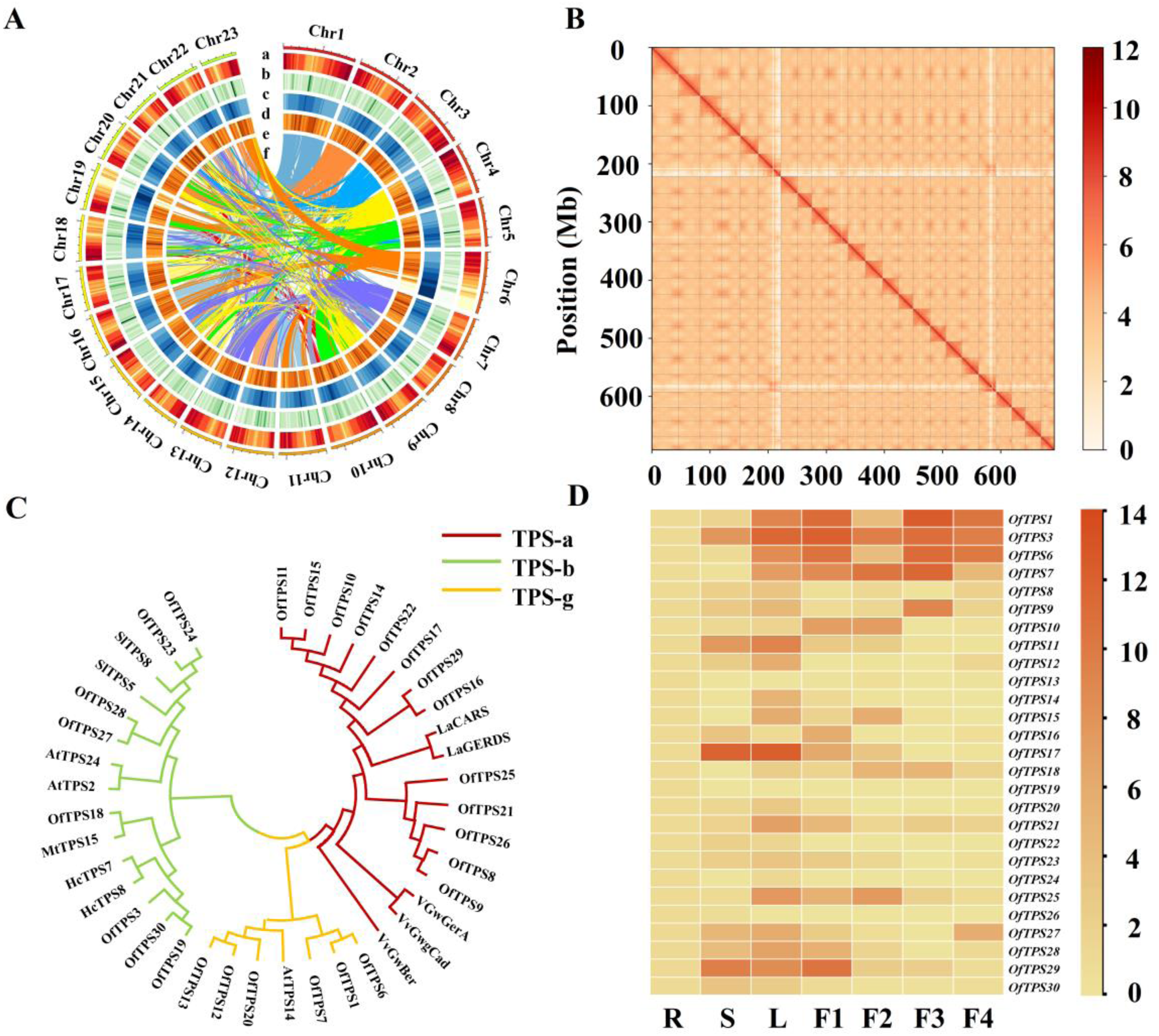
Genomic sequencing and *OfTPSs* sequence analysis of ‘BBYG’. A, Genomic characteristic map of ‘BBYG’, a: Chromosome number. b: Gene density; c: LINE; d: LTR; e: DNA transposon; f: Chromosomal collinearity; B, Hi-C interaction diagram of the genome of ‘BBYG’; C, Phylogenetic analysis of TPSs from *O. fragrans* and other plants by maximum likelihood method using MEGA7 software; D, Heatmap of *OfTPSs* expressions at different tissues root, stem, leaves and different blossoming stages (F1-F4) detected by RT-qPCR.

**Table 1.**
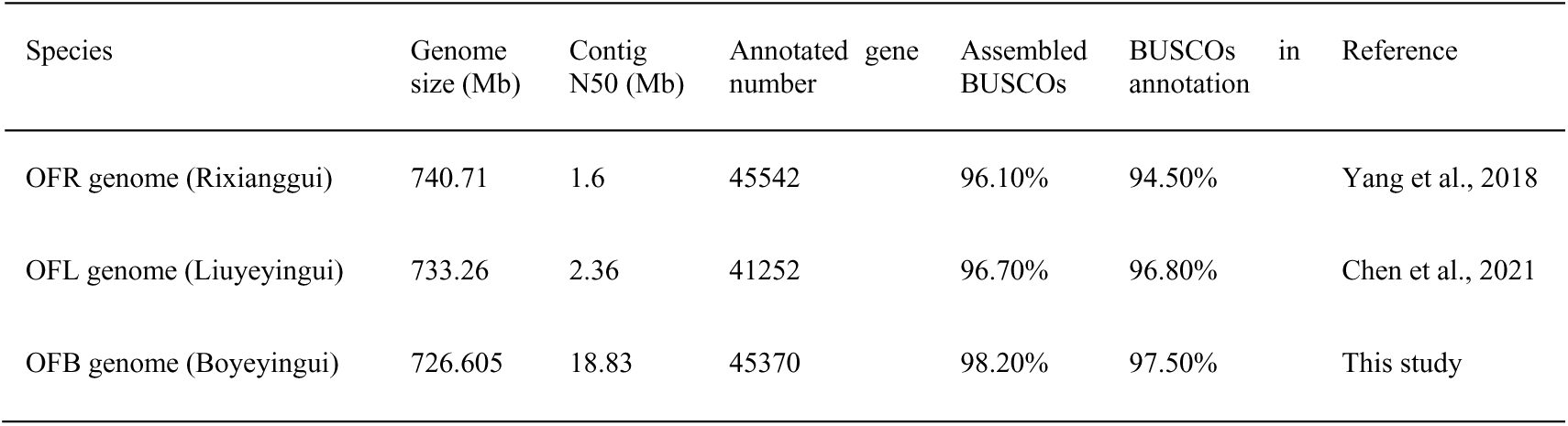
Comparisons of genome assemblies and annotations of *O. fragrans*.

Due to lack of genome sequences, only 3 *OfTPSs* were screened by NCBI blast and identified as linalool synthase genes. Here, a total of 27 *OfTPSs* were annotated, among them 25 new candidate genes were screened based on the HMM scan and BLASTp search. *OfTPS1* and *OfTPS3* were previously clarified and functionally identified as linalool synthase genes (Zeng et al., 2016). 25 full-length genes encoding putative proteins with 260 to 683 amino acids were amplified and designated as *OfTPS6-30* according to their chromosomal position (Supplementary Table 4). The majority of *OfTPSs* were found in clusters located on chromosomes I, II, V, VI, and XIX, suggesting multiple duplication and neofunctionalization events on these chromosomes (Fig. S1-2). Phylogenetic analysis classified *OfTPSs* into three subgroups, viz. TPS-a (13 members), TPS-b (7 members) and TPS-g (5 members) (Fig 2C). According to amino acid sequence alignment, *OfTPS1/6/7/12/13/20* belonging to TPS-g subfamily were typical lack of RRX_8_W, which was responsible for cyclization reactions (Fig. S3). All the other of *OfTPSs* contained all the conserved elements including RRX_8_W, DDXXD and NSE/DTE motifs that fundamentally involved in binding of Mg^2+^ or Mn^2+^ cofactors.

To compare the transcription of the MEP pathway genes with the patterns of volatile linalool during flower development, Fragments per kilobase of transcript per million mapped reads (FPKM) of putative genes were analyzed in four blossoming stages according to the transcriptome sequencings, and 37 candidate genes were involved in MEP and MVA pathway. The candidate genes in the MEP pathway had relatively higher expression levels than the candidate genes of the MVA pathway, which was in accordance with the larger amounts of the monoterpenes detected in *O. fragrans* flowers (Fig. S4, Supplementary Table 5). Reverse transcription quantitative PCR (RT-qPCR) was performed to investigate the temporal and spatial patterns of expression of all the *OfTPSs*. Six *OfTPSs* (*OfTPS1*/*3*/*6/7/9/29*) achieved high expressions in flowers compared with other tissues (Fig 2D). Notably, in agreement with the patterns of volatile linalool emission, the transcript level of *OfTPS7* substantially increased from bud to full blossoming stage and decreased afterwards (Fig 2D).

### Overexpression of *OfTPS6* and *OfTPS7* increase production of linalool both *in vitro* and in *vivo*

To verify the functions of *OfTPSs*, all the 25 new *OfTPSs* (*OfTPS6*-*OfTPS30*) were cloned and transiently transformed into tobacco leaves and *O. fragrans* flowers to determine whether they were capable of producing linalool. In the tobacco leaves, only *OfTPS6* and *OfTPS7* could synthesize linalool (Fig3A), whereas other genes did not yield any products (Fig. S5). Transient overexpression of *OfTPS6* in flowers resulted in approximately 1.61-fold increase in its transcript level and 5.35-fold increase of volatile linalool and its oxides relative to empty vector control (Fig 3C-E). Moreover, compared with the empty vector control, transient overexpression of *OfTPS7* in flowers led to approximately 3.73-fold increase in transcription level and 7.7-fold increases in volatile linalool and its oxides contents (Fig 3C-E). On the contrary, significantly lower contents of linalool and its oxides were detected in *OfTPS6* and *OfTPS7* transient silencing group compared to control group (Fig 3F-H). To further confirm the enzymatic properties of the OfTPS6 and OfTPS7 proteins, substrate specificity analyses were conducted using GPP and FPP as donors, respectively. The recombinant proteins of OfTPS6 and OfTPS7 were expressed in *Escherichia coil* and purified as soluble proteins. In vitro enzymatic assays showed that OfTPS6 and OfTPS7 were versatile enzymes with multiple products. Specifically, OfTPS6 mainly converted GPP into linalool and a few by-products such as β-myrcene, D-limonene, α-pinene and cis-β-ocimene (Fig 3B). OfTPS7 could catalyze GPP into linalool along with a small amount of β-myrcene and D-limonene (Fig 3B). OfTPS6 and OfTPS7 could not produce any volatile aroma compounds in the incubation with FPP as precursor (Fig. S6). In addition, *OfTPS7* was proved to localized in plastids where monoterpenes were produced, whereas *OfTPS6* was found to be localized in the cytosol (Fig. S7). These results clearly stated the linalool biosynthesis function of *OfTPS6* and *OfTPS7* both *in vitro* and *in vivo*.

**Figure 3.**
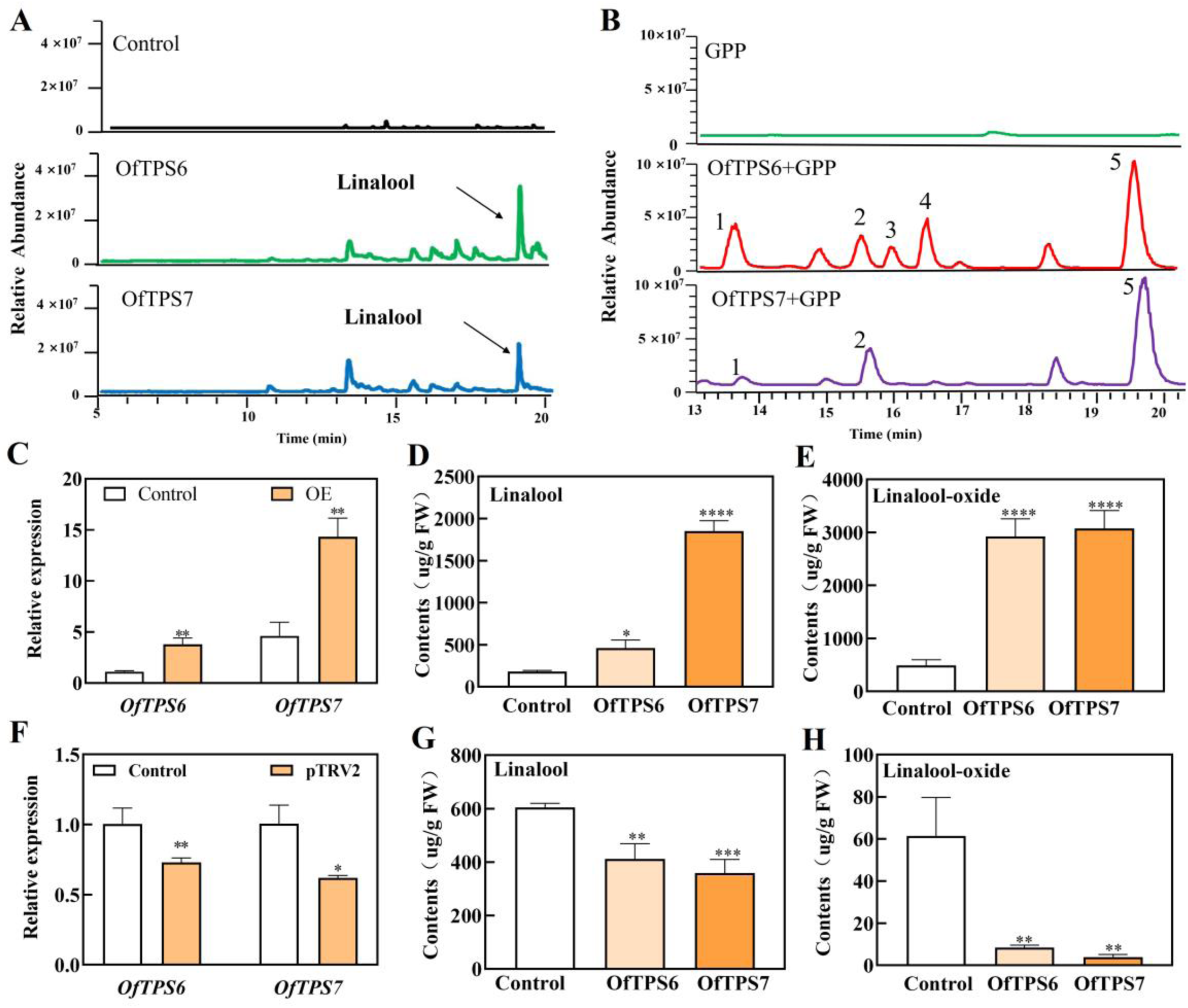
Functional validation of *OfTPS6* and *OfTPS7 in vivo* and *in vitro.* A, Aroma compound analysis of tobacco leaves of control, *OfTPS6* and *OfTPS7* overexpressed groups; B, Extracellular enzyme function of OfTPS6 and OfTPS7 (GPP as substrate), 1: β-Myrcene; 2: D-limonene; 3: α-Pinene; 4: β-Ocimene; 5: Linalool; C, Expressions of *OfTPS6* and *OfTPS7* in *O. fragrans* by RT-qPCR, Control: flowers were injected with empty vector, OE: flowers were injected with *OfTPS6* and *OfTPS7* overexpression vectors, respectively; D, Linalool contents of *O. fragrans* flowers in control, overexpressed *OfTPS6*, *OfTPS7* groups; E: Linalool oxide contents of *O. fragrans* flowers in control, overexpressed *OfTPS6*, *OfTPS7* groups; F: *OfTPS6* and *OfTPS7* expression levels in *O. fragrans* flowers in control and silencing groups; G: Linalool contents of *O. fragrans* flowers in control, silenced *OfTPS6*, *OfTPS7* groups; H: Linalool oxide contents of *O. fragrans* flowers in control, silenced *OfTPS6*, *OfTPS7* groups. Statistical significance with respect to the reference sample (control) was determined by the Student’s *t*-test. *\*P<0.05, **P<0.01, ***P<0.001*.

### *OfTPS7* plays a crucial role in linalool biosynthesis in diverse cultivars

We detected the transcription levels of all the linalool synthase genes *OfTPS1/2/5/6/7* and linalool contents in 20 cultivars to explore the key linalool synthase genes in *O. fragrans*. The results showed that *OfTPS2* exclusively expressed in two cultivars ‘BBYG’ and ‘MJYG’. *OfTPS1*, *OfTPS5 and OfTPS6* expressed in most cultivars, but their correlation coefficients with linalool content were less than 0.5. Only *OfTPS7* expressed in all 20 cultivars and its coexpression coefficient with linalool content reached the maximum value at 0.82 (Fig 4D, Supplementary Table 6-7). Additionally, spatial analysis demonstrated that *OfTPS7* achieved high expression levels at the initial and full blossoming stages in flowers, in parrel with the linalool release (Fig 4D). Therefore, we concluded that *OfTPS7* played a crucial role in linalool biosynthesis in *O. fragrans* flowers. (Fig 4E).

**Figure 4.**
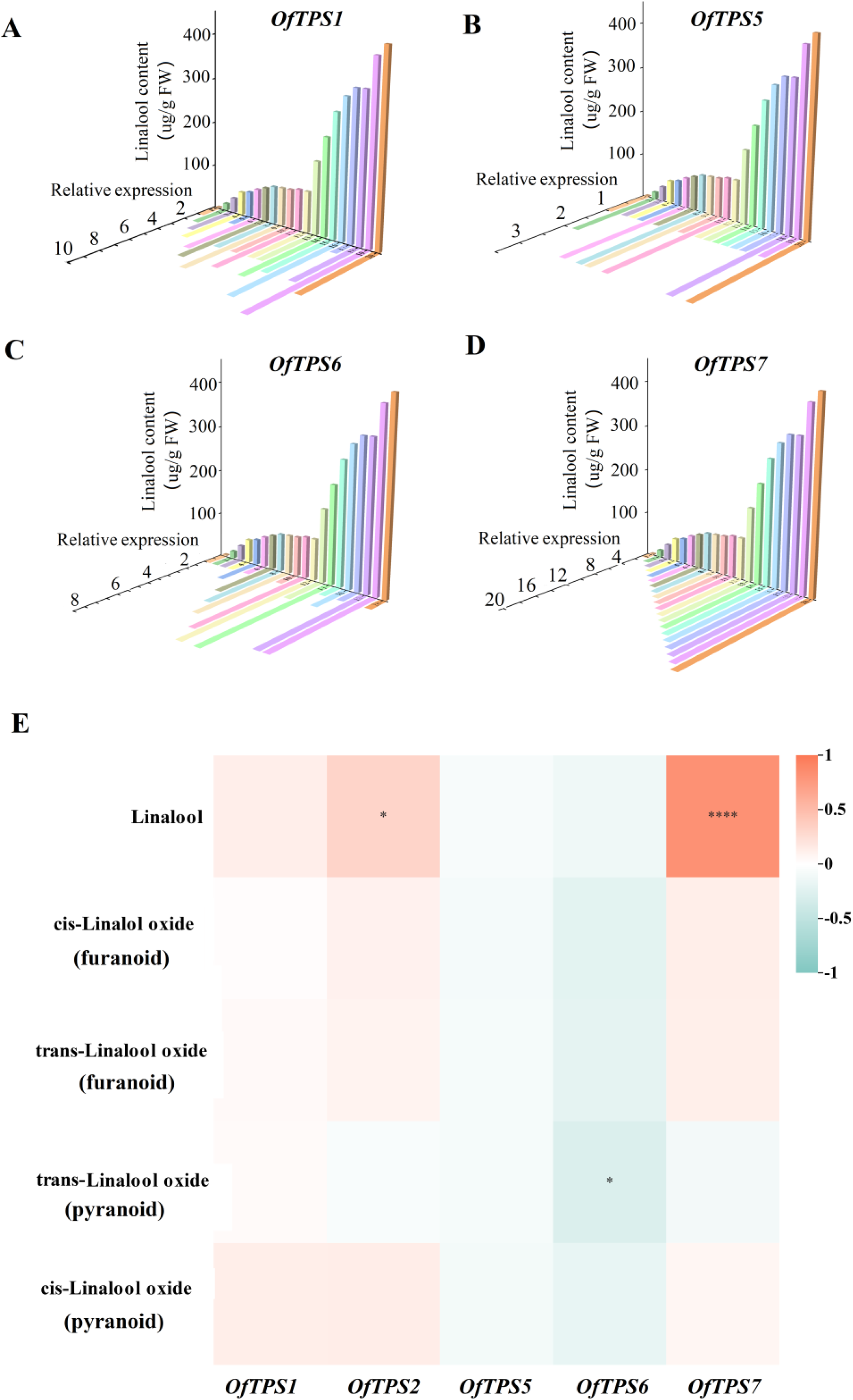
Correlation analysis of linalool synthase gene expressions with linalool and its oxides contents in 20 cultivars. A-D, Swallow-tail plot analysis of the *OfTPS1, OfTPS5, OfTPS6, OfTPS7* expression levels and linalool contents in different cultivars. E, Correlation analysis between the expression level of high expression linalool synthase gene in flowers and the content of linalool and its oxides among different cultivars.

### Y1H assay combined with WGCNA facilitate the screening of the potential transcription factor *OfWRKY33*

Having identified the crucial gene encoding the enzyme responsible for linalool formation, we next addressed the regulation of *OfTPS7*. A Y1H assay using the promoter of *OfTPS7* as the bait was applied to screen a cDNA library derived from *O. fragrans* flowers. Six putative candidates belonging to WRKY, ERF, MADS, ZAT, MYB were obtained, and one was annotated as a TF from WRKY family (Fig 5A-C, Supplementary Table 8). In order to investigate the hub genes involved in linalool biosynthesis, four important developmental stages of *O. fragrans* flowers were subjected to transcriptome sequencing in an attempt to screen potential regulators. Differentially expressed genes (DEGs) were identified by a significance threshold of log_2_ fold changes of ±1 and an adjusted P value<0.01. A total of 19199 non-redundant DEGs were retained for WGCNA after filtering, leading to four enriched modules related to linalool and other terpenoids, respectively. Analysis of module-trait correlations revealed that the blue module genes containing 1890 TFs were closely associated with the synthesis of linalool (*r*=0.66, *P*=0.02) (Fig. 5D). In addition, DEGs in blue module were analyzed by a Venn diagram to screen differentially expressed genes in four blossoming stages (Fig. S8AB). Among them, the correlation coefficients of 6 TFs with *OfTPS7* were higher than 0.85 and RT-qPCR results further confirmed that the expression patterns of 4 TFs were generally parallel with *OfTPS7*. Spatial and temporal analysis revealed that *OfWRKY33* not only achieved higher transcript level in flowers than in stem, leaf and root, but also performed similar trend with linalool release in flowers (Fig. 6AB). Meanwhile, Y1H assay also showed potential interaction between *OfWRKY33* and *OfTPS7* (Fig. 5D). These data suggested that *OfWRKY33* might be connected with linalool biosynthesis (Fig 5-EF).

**Figure 5.**
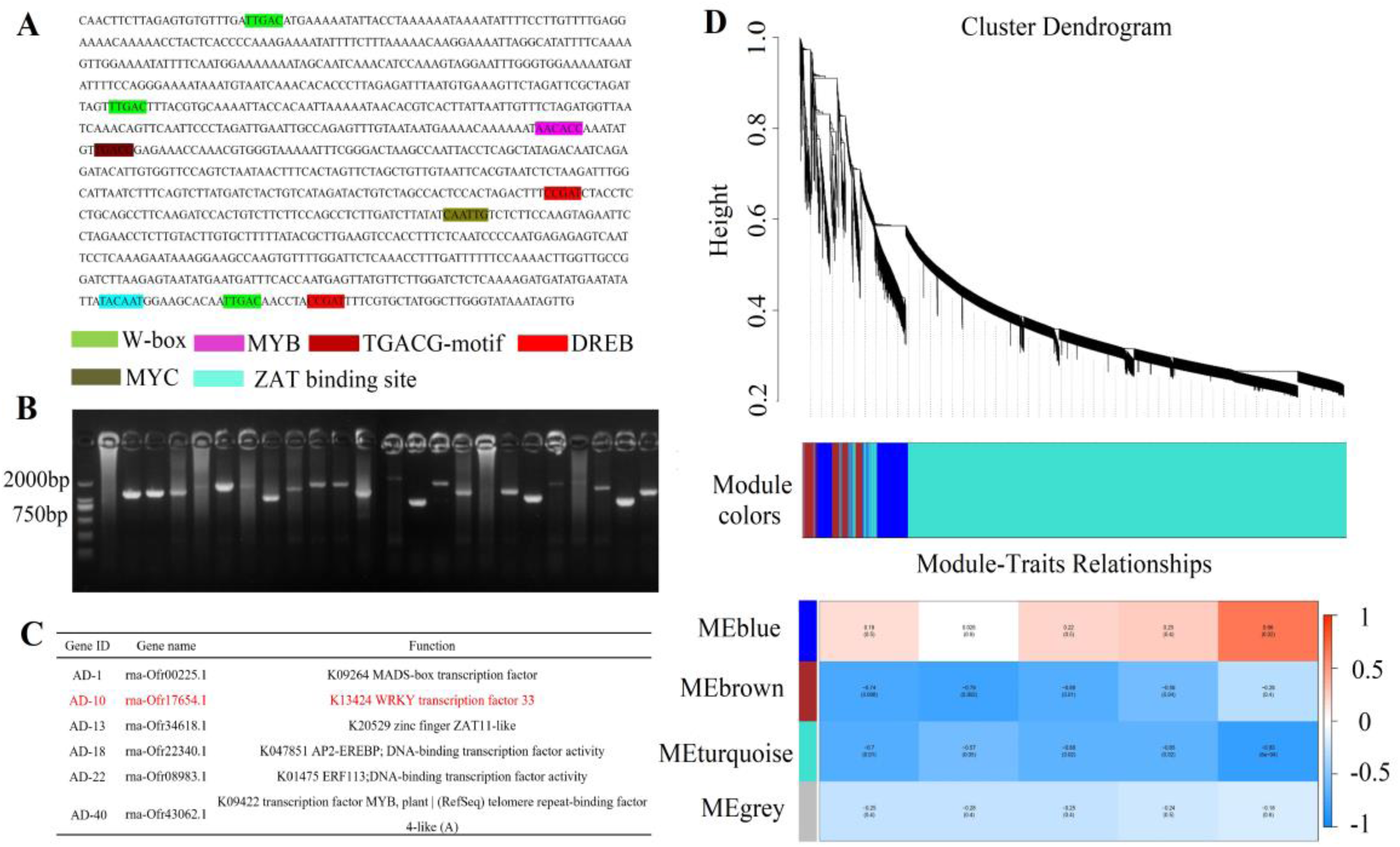
Screen of potential regulators of OfTPS7 combined by Y1H and WGCNA. A, Analysis of cis acting elements in *OfTPS7* promoter; B, Detection of positive bacteria in yeast single impurity screening library; C, Functional annotation of yeast single impurity screening library positive bacteria; D, WGCNA analysis of transcriptome at different flowering stages of ‘BBYG’.

**Figure 6.**
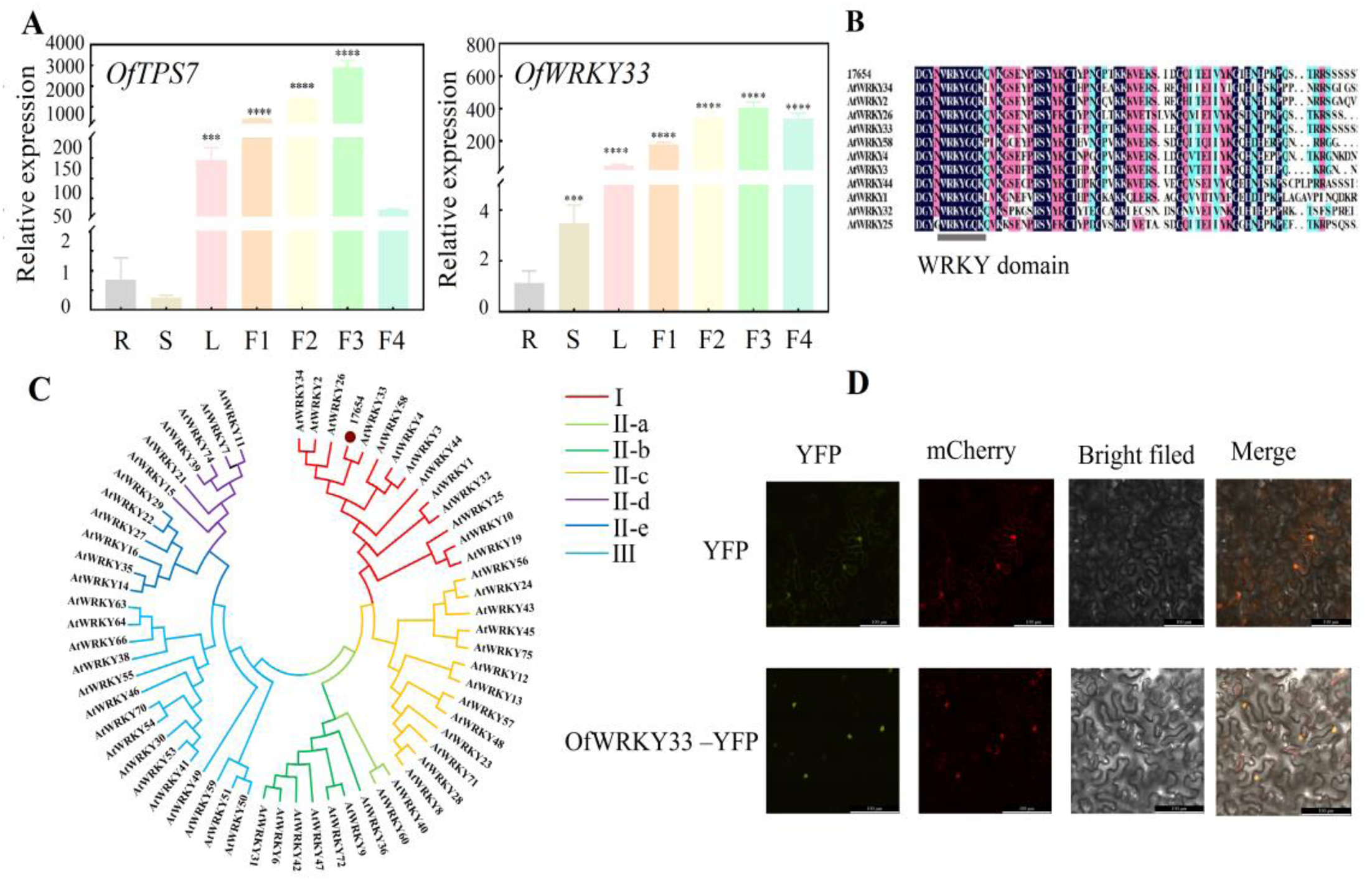
Expression patterns, sequence analysis and subcellular localization of *OfWRKY33*. A, Expression levels of *OfTPS7* and *OfWRKY33* in different tissue parts and flowering stages of ‘BBYG’; B, Sequence alignment between OfWRKY33 and AtWRKYs; C, Phylogenetic analysis of WRKY proteins from *O. fragrans* and other plants. The phylogenetic tree was constructed by using the maximum likelihood method through MEGAX software. Detailed information of TPS proteins was provided in Supplementary Table 10; D, *OfWRKY33* subcellular localization, YFP fluorescence indicated the location of each fusion protein (shown in yellow), the location of nucleus was determined by nuclear localization marker (shown in red). Column labelled ‘Merged’ represented all the combined fluorescent signals. Bars, 100 μm. Statistical significance with respect to the reference sample (control) was determined by the Student’s *t*-test. *\*P<0.05, **P<0.01, ***P<0.001*.

### OfWRKY33 is a nucleus-located transcription factor

The *OfWRKY33* showed 50% amino acid similarity with *AtWRKY33* from *A. thaliana* (GenBank accession number NP_181381.2) (Fig 6C). The full-length cDNA sequence of *OfWRKY33* (GenBank accession PP598872 and *O. fragrans* genome accession number Ofr17654) contained an open reading frame of 1468 bp, encoding a polypeptide of 489 amino acid residues with a conserved WRKYGQK domain near its C terminus (Fig 6D). *OfWRKY33* had a calculated molecular mass of 53.796kDa. Multiple sequence alignment indicated that *OfWRKY33* was clustered into WRKY-I subfamily. To determine the subcellular localization, a *CaMV35S*:*OfWRKY33*-*YFP* construct and a *CaMV35S*: *YFP* control construct were introduced into *Nicotiana benthamiana* leaves, respectively. Microscopy illustrated that *OfWRKY33-YFP* was located exclusively in the nucleus and overlapped with the mCherry, whereas *CaMV35S*: *YFP* was distributed evenly throughout the cell (Fig 5E).

### *OfWRKY33* enhances the linalool content and affects the transcript of some MEP pathway genes in transgenic flowers

To further verify the roles of *OfWRKY33* in MEP pathway, we applied transiently transgenic experiments in *O. fragrans* flowers. *Agrobacterium tumefaciens* GV3101 containing the plasmid *PK7WG2D-OfWRKY33* and the viral-induced gene silencing (VIGS) plasmid *pTRV2*-*OfWRKY33* vectors were vacuum infiltrated into *O. fragrans* flowers at the initial blossoming stage, respectively. The expressions of *OfWRKY33* and *OfTPS7* were significantly elevated in the overexpressed flowers, resulting in remarkable promotion of linalool and its oxides contents (Fig 6A, C). The transcript levels of MEP pathway genes were also monitored. Notably, the expressions of *OfDXS1*, *OfDXR2*, and *OfIDI2*, which had higher FPKM values in the transcriptome of different flowering stages of ‘BBYG’, were also stimulated in *OfTPS7* over-expressed flowers (Fig 7 B, E, Fig. S4). The transcript levels of other linalool-related genes namely *OfDXS5*, *OfGPPS1* and *OfGPPS5*, did not show similar patterns in the *PK7WG2D*-*OfWRKY33* flowers. *OfWRKY33*-silenced flowers contained both remarkably lower gene expressions (especially *OfDXS1*, *OfGPPS1*, and *OfGPPS5*) and linalool contents compared with the control group (7.41-fold and 5.85-fold) (Fig7 D, F). These results indicated that *OfWRKY33* was a positive modulator of linalool biosynthesis.

**Figure 7.**
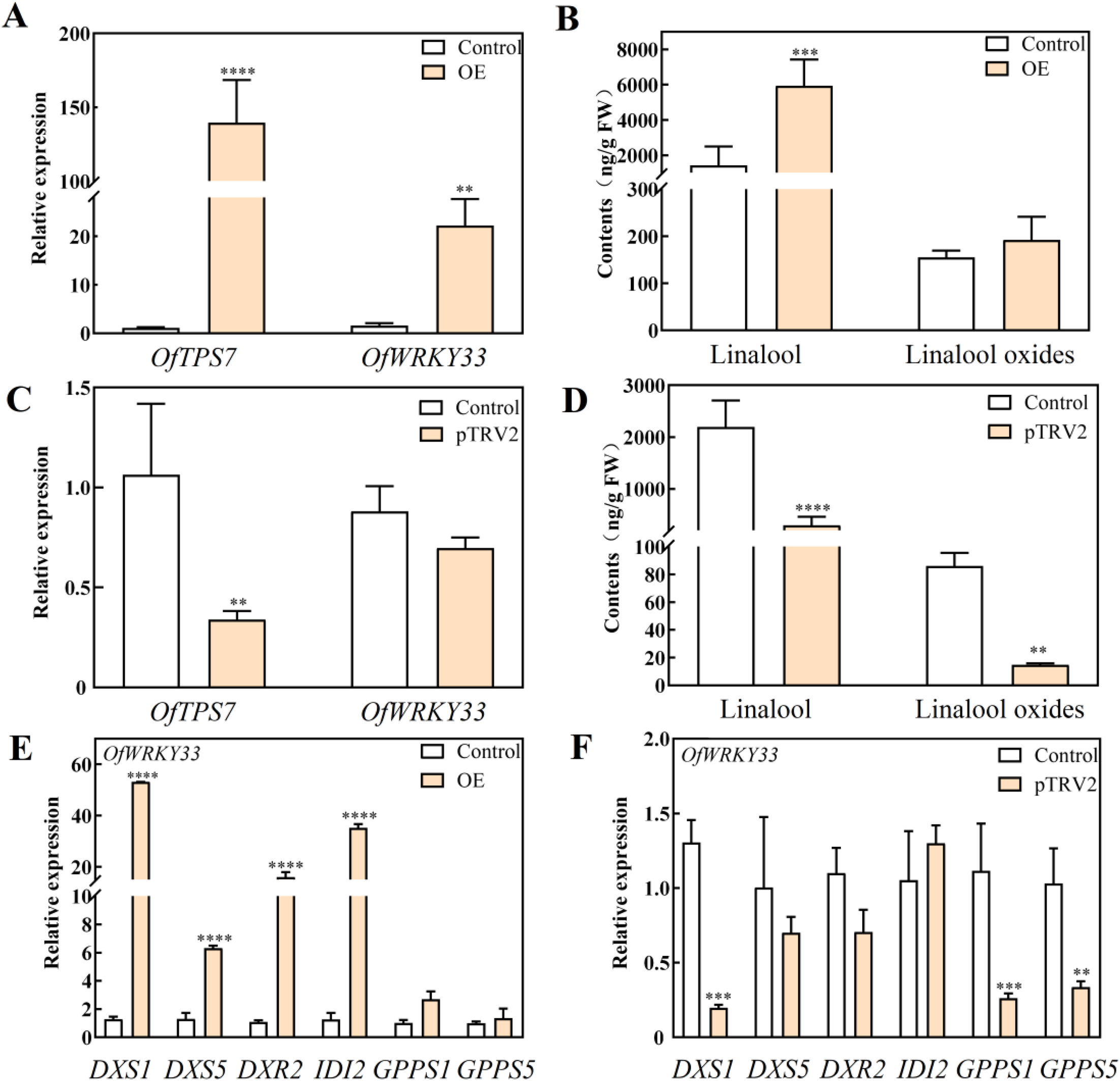
Functional verification of *OfWRKY33* in *O. fragrans* flowers. A, Gene expression level of *OfWRKY33* and *OfTPS7* in overexpressing strain; B, Linalool and linalool oxide content in overexpressed strains of *OfWRKY33*; C, Gene expression level of *OfWRKY33* and *OfTPS7* in silenced strain; D, *OfWRKY33* silent line linalool and linalool oxide content; E, Expression analysis of important genes in MEP pathway in overexpressed and control strains of *OfWRKY33*; F, Expression analysis of important genes in MEP pathway in silenced and control strains of *OfWRKY33*. Statistical significance with respect to the reference sample (control) was determined by the Student’s *t*-test. *\*P<0.05, **P<0.01, ***P<0.001*.

**Figure 8.**
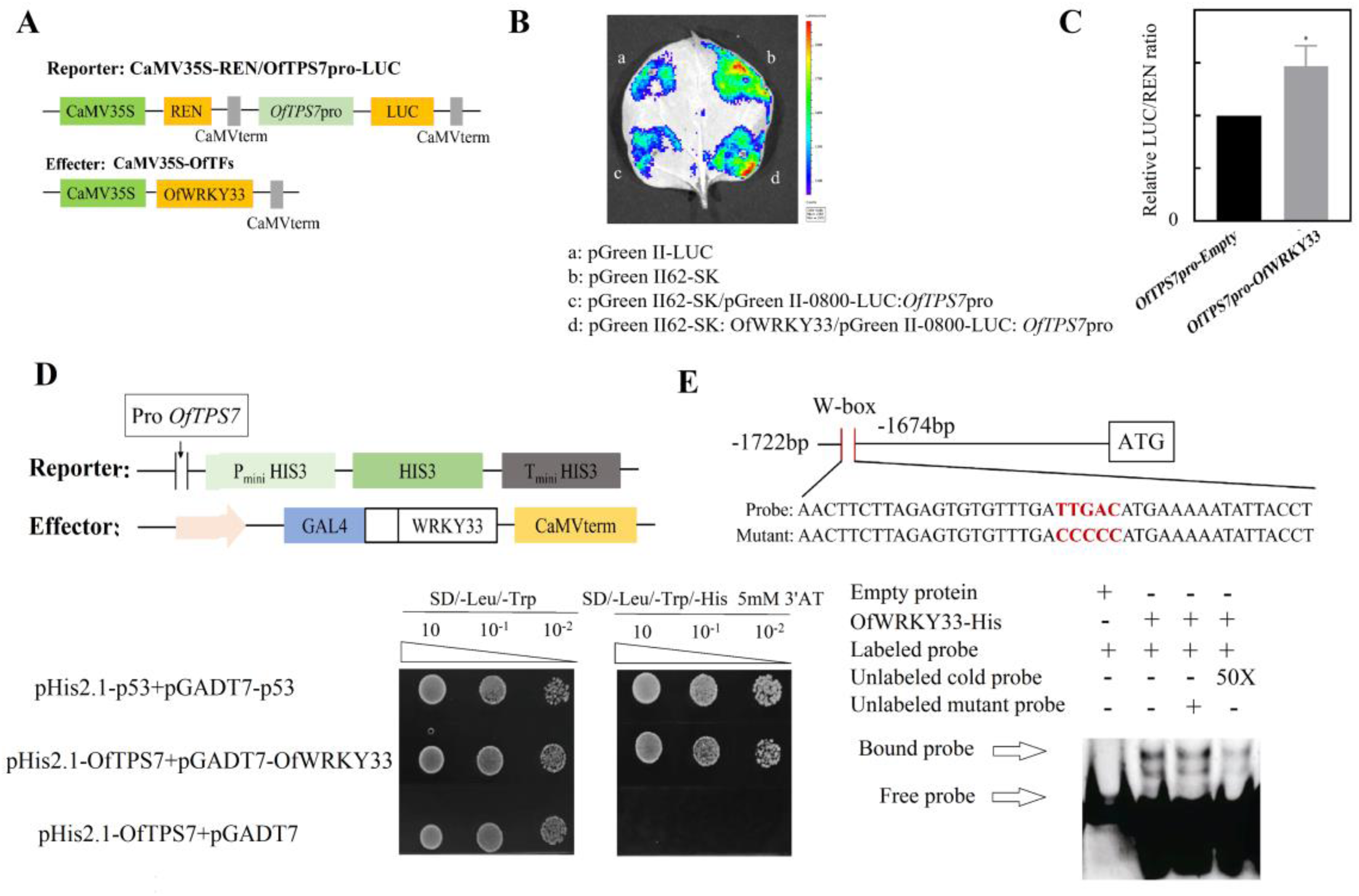
*OfWRKY33* binds directly to the W-box in the *OfTPS7* promoter sequence. A, Schematic diagram of dual luciferase carrier, the effector vector *OfWRKY33* was constructed on the pGreen-62-SK vector driven by CaMV35S, and the reporter vector *OfTPS7* promoter was constructed on the pGreen II-0800 LUC vector to drive the expression of LUC; B, The transient fluorescence activity of tobacco indicates that *OfWRKY33* has transcriptional activation activity and can promote the expression of *OfTPS7*; C, Relative fluorescence activity of *OfWRKY33* transcriptional activity; D, Yeast single heterozygous display of interaction between *OfWRKY33* and *OfTPS7* promoter; E, OfWRKY33 binds directly to the *OfTPS7* promoter, EMSA probe, Mut represents mutation probe, and mutation sites are represented by black dashed boxes, with “+” and “-” indicating the addition or absence of probes or proteins; 50× represents 50 fold cold probe, arrow indicating protein DNA complex or free probe position. The error line represents the standard deviation of 12 biological replicates. Statistical significance with respect to the reference sample (control) was determined by the Student’s *t*-test. *\*P<0.05, **P<0.01, ***P<0.001*.

**Figure 9.**
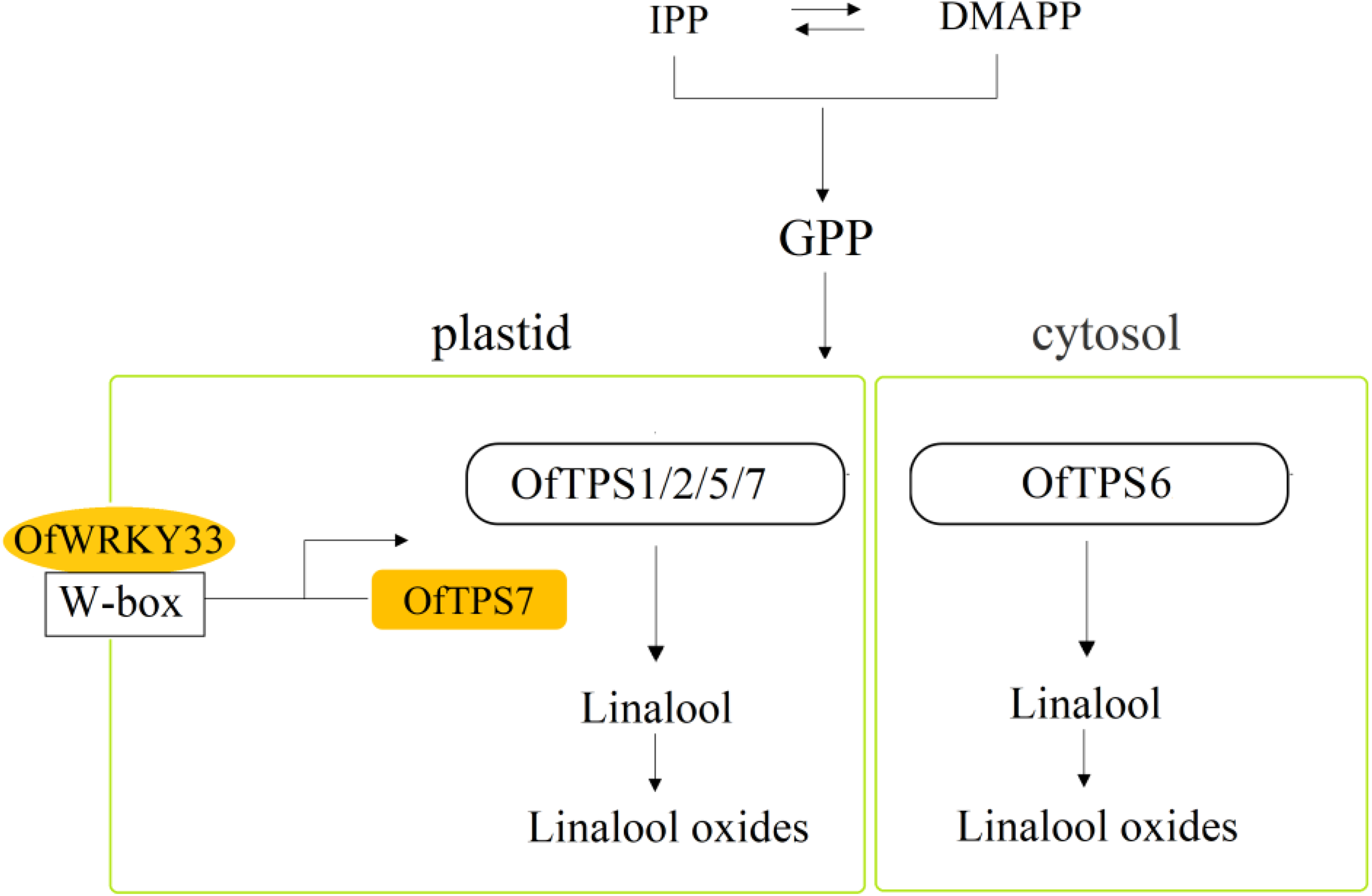
The linalool biosynthesis and its transcriptional regulation in *O. fragrans* flowers. GPP serve as the donor, *OfTPS1/2/5/7* are located in plastid and express in 20 cultivars, whereas *OfTPS6* is located in cytosol and express in 2 cultivars. Among them, *OfTPS7* is a key linalool synthase gene and OfWRKY33 can directly bind to the W-box of *OfTPS7* promoter to manipulate linalool formation.

### *OfWRKY33* directly binds to the promoters of *OfTPS7* and activates its transcription

To further determine whether *OfWRKY33* could activate the expression of *OfTPS7*, we first performed a dual LUC assay. The promoter of *OfTPS7* was inserted into LUC reporter vector, and the coding DNA sequence (CDS) of *OfWRKY33* was cloned and inserted into the effector vector (Fig 7A). The results showed that compared with the empty vector, *OfWRKY33* significantly elevated the relative LUC expression (LUC/LEN ratio) driven by the *OfTPS7* promoter (Fig 7C). Furthermore, LUC imaging assays also confirmed that *OfWRKY33* activated the *OfTPS7* promoter *in vivo*. The co expression region (*OfWRKY33*+*OfTPS7pro*-Luc) exhibited a strong fluorescent signal, whereas the control region (EV+*OfTPS7pro*-Luc) displayed a weak fluorescent signal (Fig7B). *OfWRKY33* also displayed strong interactions with the promoters of *OfTPS7* in Y1H assays (Fig7D). LUC and Y1H experiments were also conducted to test the relationship between *OfWRKY33* and *OfDXS1*, *OfDXR2* and *OfIDI2*, but no direct interaction was observed (Fig. S9-10).

Electrophoretic mobility shift assays (EMSA) were subsequently performed to test whether the OfWRKY33 protein bound to the promoter of *OfTPS7*. A specific shifted band was observed when the recombinant protein OfWRKY33-His and a labeled probe coexisted and that His shifted band was weakened by the unlabeled probe, whereas it was not weakened by the mutated unlabeled probe (Fig 7F). These results suggested that the OfWRKY33 protein specifically bound to the W-box on the *OfTPS7* promoter *in vitro*.

## Discussion

Terpenoids are abundant aroma compounds in different flowers and linalool along with its oxides are determined as vital aroma active compounds in various plants (Yu et al., 2020; Li et al., 2022; Wei et al., 2021). Although significant progress has been made in identifying the biosynthetic pathways of linalool in many species, the hub linalool synthase genes together with their underlying transcriptional regulation is not well understood. Here, we elucidated the dominant role of *OfTPS7* in the synthesis of linalool in multiple cultivars and deciphered the novel OfWRKY33-driven regulatory network of linalool biosynthesis in *O. fragrans* flowers. We confirmed that OfWRKY33 acted as a positive regulator of linalool formation by directly binding to the *OfTPS7* promoter and activating the expression of *OfTPS7* together with other key pathway genes.

### Comprehensive study of *OfTPSs* provides global view of terpenoid formation

TPSs are the metabolic gatekeepers in the biosynthesis of diverse terpenoids in plants. TPS gene family is a mid-size family of highly diversified sequences and functions, with gene numbers ranging from approximately 20 to 180 in the plant genomes (Shang et al., 2020; Yu et al., 2021; Zhou et al., 2022; Jiang et al., 2024). The first linalool synthase gene (*LIS*) from the flowers of *Clarkia breweri* was isolated in 1996 (Dudareva et al., 1996). In recent years, molecular mechanism study of terpenoids is developing rapidly, the characterization of TPSs have been achieved in various plants such as *Rosa hybrida* (Chen et al., 2021; Corentin et al., 2023), *Fressia× hybrida* (Gao et al., 2018), *Lathyrus odoratus* (Bao et al., 2020), *Dendrobium officinale* (Yu et al., 2020), *Chrysanthemum indicum* (Zhou et al., 2021) and so on. In fact, only a small proportion of TPSs in genome are responsible for yielding terpenoids in plants. For instance, 2 members out of 40 *AtTPSs* were capable of forming over 20 sesquiterpenes in *A. thaliana* (Tholl et al., 2005). To date, all the *OfTPSs* annotated in the genome have not been thoroughly characterized to gain a comprehensive view of terpenoid formation in *O. fragrans*. Five linalool synthase genes *OfTPS1/2/5/6/7,* belonging to TPS-g subfamily, were identified as linalool synthase genes. *OfTPS3/4* were clustered into TPS-b subfamily. *OfTPS3* was capable of producing trans-β-ocimene, while *OfTPS4* could produce the sesquiterpene α-farnesene (Zeng et al., 2016).

The composition of terpenoids was closely related to subcellular localization, gene function and gene expression (Zhou and Pichersky, 2020; Li et al., 2024). *OfTPS7* exclusively distributed in plastid, where GPP was dominantly produced as substrate for monoterpenes. However, *OfTPS6* was found to be localized in the cytosol (Supplementary Fig. S7), where a proportion of GPP might originate via crosstalk from MEP pathway to MVA pathway (Conart et al., 2023). In addition, *OfTPS7* presented relatively high expression and was closely related with the linalool content in 20 cultivars (Fig 4). However, *OfTPS2* was expressed in 2 cultivars. The correlation coefficient of *OfTPS1/2/5/6* expression with linalool contents were lower than 0.5. Thus, we proposed that although *OfTPS1/2/5/6/7* produced the overlapping linalool, *OfTPS7* was the core linalool synthase gene in *O. fragrans* flowers.

Although OfTPS6/7 could give rise to multiple monoterpenes with incubation of GPP, they could only produce linalool in planta. In this case, the loss of catalytic abilities of *OfTPS6/7* might be a consequence of the important biological roles of linalool and its derivatives for *O. fragrans.* Moreover, linalool tended to be abundantly hydroxylated and glycosylated in the late blossoming stage possibly acting as anti-microbial and anti-herbivorous defense for the forthcoming fruit (Zheng et al., 2019; Dong et al., 2024).

### Characterization of OfTPSs lays foundation for future metabolic engineering

Terpenoids such as linalool and its oxides, α-ionone, β-ionone are extremely abundant in *O. fragrans* flowers. They have been widely applied as ingredients in food, cosmetics, therapeutic and flavoring products (Jiang et al., 2023). Conventionally, these aroma compounds are obtained from plant extracts using organic solvents or distillation. But the great challenges for industrial utilization of *O. fragrans* flowers are the short blossoming period, postharvest treatment and essential oil extraction technology. It is extremely costly and laborious if high-quality products are required. Thus, microbial biosythesis has emerged as alternative source with great promise for meeting the increasing demand for terpenoid biomanufacturing (Zhu et al., 2021). The characterization of functional OfTPSs provides useful gene resources for metabolic engineering in *Saccharomyces cerevisiae* and *Yarrowia lipolytica*.

### Transcriptional regulation expands understanding of molecular mechanism of linalool formation

As the terminal biosynthesis gene in terpenoid biosynthesis, *TPS* was the hot topic of aroma compound formation. However, only a limited number of TFs were reported to be directly involved in the regulation of TPSs. For instance, *LaMYC7* directly bound to the promoter of *LaTPS76* to affect the formation of linalool and caryophyllene biosynthesis in *Lavendula angustifolia* (Dong et al., 2024). *SlMYB75* was capable of binding to the *SlTPS12*, *SlTPS31* and *SlTPS35* and inhibiting their transcription, thereby negatively affecting the biosynthesis of β-caryophellene and α-humulene in *Solanum lycopersicum* (Gong et al., 2021). In flowers of *Freesia hybrid*, FhMYB21L2 activated transcription of linalool synthase gene *FhTPS1* by binding to the MYBCORE sites in the promoter (Yang et al., 2020). Progress in characterization of upstream TFs influencing terpenoids lags far behind compared with color-related chemicals such as anthocyanin and carotenoid.

Since the functions of *OfTPSs* have been thoroughly elucidated and *OfTPS7* was proved to be closely associated with the linalool formation, the transcriptional regulation has become a new focus in *O. fragrans*. *OfMYB21* was previously proved to directly bind to *OfTPS2* promoter and promote linalool biosynthesis in *O. fragrans* flowers (Lan et al., 2023). However, our study demonstrated that *OfTPS2* could only be detected in 2 cultivars, therefore the regulation role of *OfMYC2* might be limited. In the current study, WGCNA derived from transcriptomes, yeast single impurity screening library, expression analysis and functional identification were integrated to claim that *OfWRKY33* could positively regulate linalool biosynthesis. A variety of approaches (i.e. Y1H, EMSA and dual-luciferase assay) were further conducted to unravel that OfWRKY33 bound directly to a specific W-box in the promoter of *OfTPS7.* The above results provided evidence that OfWRKY33 acted as a direct positive regulator in linalool biosynthesis.

OfWRKY33 also had influenced multiple steps throughout the MEP pathway. The expression patterns of *OfDXS1*, *OfDXR2*, and *OfIDI2* were consistent with *OfWRKY33* in the over-expression and silenced *O. fragrans* flowers. Although these were not housekeeping genes, they were constitutively and highly expressed in *O. fragrans* flowers to provide substrate for linalool formation. Similarly, *SlSCL3* simultaneously stimulated the expressions of *SlTPS* along with upstream isoprenoid precursor pathway genes including *SlHMGS*, *SlHMGR* and *SlDXS*, which determined the flux through this pathway in *S. lycopersicum* (Yang et al., 2021). However, no binding activity was detected between *OfWRKY33* and the promoters of *OfDXS1*, *OfDXR2*, and *OfIDI2* (Supplementary Fig. S9), suggesting that activation of transcription might require the presence of other proteins. TFs can form a complex to cooperatively mediate terpenoid biosynthesis (Yang et al., 2020; Gong et al., 2021). The physical interactions between TFs affected the protein-DNA interactions therefore influencing the gene expression. For example, protein complex of ZmMYC2-ZmEREB92 exhibited stronger DNA binding ability to *ZmTPS6* than ZmMYC2 alone (Fu et al., 2023). Future studies should investigate the synergistical regulation concerning the interaction of *OfWRKY33* with other TFs, which will expand our understanding of the regulatory role. WRKYs are one of the largest family of plant-specific TFs, they play pivotal roles in plant defense, growth and development, morphogenesis of trichomes and embryos, hormone signaling and secondary metabolite biosynthesis (Zhou et al 2020; Chen et al 2024). Our study proposed that *OfWRKY33* directly interacted with functional genes to regulate the aroma compound biosynthesis, providing new insights for research of WRKY.

In summary, we first characterized the key linalool synthase gene *OfTPS7* in *O. fragrans* flowers. In addition, we provide a schematic of a global view of transcriptional regulation of *OfTPS7*. *OfWRKY33* could directly bind to the W-box of *OfTPS7* promoter and simultaneously activated multiple MEP pathway genes. These findings expand our understanding in the complex molecular mechanism of linalool biosynthesis in *O. fragrans* flowers.

## Material and methods

### Plant materials

The flowers of ‘BBYG’ were collected at different blossoming stages, viz. bud stage (F1), initial blossoming stage (F2), full blossoming stage (F3), and late blossoming stage (F4) in Xianning (28°32′N, 110°36′E), Hubei Province, China. After volatile aroma compound sampling, the floral samples were promptly frozen in liquid nitrogen and stored at −80℃ for further use. All the above materials were sampled with 3 biological replicates. *N. benthamiana* plants were grown in a glasshouse under a 16h: 8h, light: dark cycle and at 24℃ in 60% humidity and under 250 μmol m^−2^ s^−1^ intense luminosity.

### Genome sequencing of *O. fragrans* ‘BBYG’

We used a total of high-quality 15 μg genomic DNA derived from four stages of flowers to construct a SMRTbell target size library using the SMRTbell Express Template Prep kit 2.0 (Pacific Biosciences) for PacBio-HiFi sequencing. We sheared genomic DNA to the expected size of fragments for sequencing on a PacBio Sequel II (Pacific Biosciences, Menlo Park, CA, USA) Binding Kit 2.0 in Frasergen Bioinformatics Co. Ltd. (Wuhan, China). Hi-C libraries were constructed according to previous studies (Dudchenko et al., 2017). The Hi-C libraries were quantified and sequenced on the MGI-seq platform (BGI, China).

### Genome assembly and annotation

All subread data from SMRT sequencing were used for ‘BYYG’ genome assembly. The draft genome assembly was obtained using mecat 2 with the default parameters. The gcpp in the SMRT link toolkit was performed to correct errors after the initial assembly of the genome (Roach et al., 2018). Next, SMRTbell libraries were sequenced on a PacBio Sequel II system and consensus reads (HiFi reads) were generated using ccs software (https://github.com/pacificbiosciences/unanimity) with the parameter ‘-minPasses 3’. To further validate and improve the assemblies, we generated 91.3× PacBio HiFi reads for the three accessions. These long (∼15 kb) and highly accurate (>99%) HiFi reads were assembled using hifiasm v. 0.14-r312 with default parameters, and the gfatools (https://github.com/lh3/gfatools) was used to convert sequence graphs in the GFA to FASTA format (Driguez et al., 2021). Finally, for anchored contigs, clean reads were generated from the Hi-C library and were mapped to the preliminary assembly using Juicer with default parameters. Paired reads mapped to different contigs were used for the Hi-C associated scaffolding. Self-ligated, non-ligated, and other invalid reads were filtered out. We applied 3D-DNA to order and orient the clustered contigs. Then, Juicer was used to filter the sequences and cluster them, and the Juicebox was applied to adjust chromosome construction manually. We finally anchored the scaffolds on twenty-three chromosomes. In addition, the BUSCO pipeline was used to assess the completeness and accuracy of the ‘BYYG’ genomes (Durand et al., 2016). The two methods are combined to identify the repeat contents in our genome, homology based and de novo prediction. We predicted protein-coding genes of the ‘BYYG’ genome using three methods, including abinitio gene prediction, homology-based gene prediction and RNA-Seq-aided gene prediction. Gene functions were inferred according to the best match of the alignments to the National Center for Biotechnology Information (NCBI) Non-Redundant (NR), TrEMBL, InterPro and Swiss-Prot protein databases using BLASTP (ncbi blast v2.6.0+) and the Kyoto Encyclopedia of Genes and Genomes (KEGG) database with an E-value threshold of 1E-5. The protein domains were annotated using PfamScan (pfamscan_version) and InterProScan (v5.35-74.0) based on InterPro protein databases. The motifs and domains within gene models were identified by PFAM databases. Gene Ontology (GO) IDs for each gene were obtained from Blast2GO. In total, approximately 43612 (about 63.13%) of the predicted protein-coding genes of ‘BYYG’ could be functionally annotated with known genes, conserved domains, and Gene Ontology terms.

### Transcriptome sequencing of flowers in four blossoming stages

In order to obtain effective information for gene annotation, the Iso Seq method was performed using the SMRT sequencing platform to generate full-length transcripts. The RNA from flowers at four different blooming stages were prepared from the same tree and processed for library construction. Total RNA was extracted using TRIzol reagent (Invitro-gen) according to the manufacturer’s protocol. A total of 12 RNA-seq libraries (three biological replicates at four blossoming stages) were prepared using the Clontech SMARTer cDNA synthesis kit according to the manufacturer’s recommendations and were then sequenced on the PacBio Sequel II platform at Frasergen Bioinformatics Co. Ltd.

### Identification of differentially expressed genes and co-expression network modules

After discarding undetectable or relative low expression genes (Transcripts Per Million reads, TPM<10), DEGs (coefficient of variation, CV > 0.5) were extracted to generate co-expression network modules by weighted gene co-expression network analysis (WGNCA) package. Benjamini-Hochberg correction (Benjamini and Hochberg, 1995) was used to adjust the original P-value in Baggerly’s test to minimize the false discovery rate (FDR). A DEG was declared if the associated P_FDR_<0.05 was observed. WGCNA network construction and module detection was conducted using an unsigned type of topological overlap matrix (TOM), a power β of 10, a minimal module size of 30, and a branch merge cut height of 0.25. The module eigengene (the first principal component of a given module) value was calculated to evaluate the association of modules and terpenoid contents in the 12 samples.

### Gene isolation, phylogenetic tree construction and multiple sequence alignment

We used HMMER software to search for *OfTPSs* and TFs in the genome. *OfTPSs* and TFs containing conserved domains were subjected to the online web tool GSDS2.0 for gene structure analysis. The full-length cDNAs of *OfTPSs* and TFs were obtained by PCR amplification with gene-specific primers (Supplementary Table 9) based on annotated results from the genome database. The cDNA used as template was synthesized from RNA derived from four blossoming stages flowers. Multiple full-length amino acid sequence alignments of TPS and WRKY proteins were performed using DNAMAN software. The phylogenetic trees were constructed using maximum likelihood method in MEGA7.0 software (Kumar et al., 2016). Bootstrap analysis was performed using 1000 replicates to evaluate the reliability of different phylogenetic group assignments.

### RNA extraction and RT-qPCR

Total RNA was extracted from the samples using a kit (Aidlab Biotechnology, Beijing, China). cDNA was synthesized using Hiscript II QRT SuperMix for qPCR. Three biological replicates were tested for all samples using *β-actin* as the endogenous control gene for data normalization. RT-qPCR was conducted using Roche Lightcycle 480 system (Roche) and relative expression levels were calculated by the 2^−△△Ct^ method (Livak and Schmittgen, 2001). The RT-qPCR primers used in this study are listed in Supplementary Table S9.

### Transient transformation of *N. benthamiana* leaves and *O. fragrans* flowers

All the coding regions of newly found *OfTPSs* derived from the ‘BYYG’ genome were amplified and the PK7WG2D-OfTPSs plasmids were transferred into *Agrobacterium* GV3101 and cultivated until their OD_600_ values reaching 0.6-0.8. The transient transformation of *OfTPSs* into the *N. benthamiana* leaves was performed as previously described (Gao et al., 2018).

PK7WG2D-OfTPS6, PK7WG2D-OfTPS7, PK7WG2D-GFP, pTRV2-OfTPS6, pTRV2-OfTPS7 and pTRV1 plasmids were constructed and transferred into *Agrobacterium* GV3101, respectively. Freshly grown *Agrobacterium* cultures reaching an OD_600_ of 0.6-0.8 were centrifuged and resuspended in infiltration media [10 mM 2-(n-Morphorinic) Ethanesulfonic acid, 10 mM MgCl_2_, 200 μM acetosyringone] and incubated without shaking at room temperature for 2-3 h. *O. fragrans* flowers at the initial blossoming stages were collected for vacuum infiltration at 0.08 Mpa for 5 min. The residual *Agrobacterium* solutions were removed with sterile water and the flowers were maintained in 5% sterile sucrose solution in the dark for 60 h. After incubation, RT-qPCR and GC-MS assays were carried out to determine the gene expression and aroma compound contents. The functional identification of OfWRKY33 were also conducted using the above methods.

### Heterologous expression and in vitro enzyme activity in *Escherichia coli*

TargetP 2.0 (https://services.healthtech.dtu.dk/services/TargetP-2.0/) and WoLF PSORT (Horton et al., 2007) were applied to predict the signal peptides of *OfTPS6* and *OfTPS7*. *OfTPS6* contained a segment of signal peptide. The truncated *OfTPS6* and *OfTPS7* coding region sequences on the Pet21b vectors were constructed for removal of signal peptide. The vectors were then transformed into Rosetta2 (DE3) cells for further induction and cultivation for 14-16 hours (Gao et al., 2018). Recombinant proteins were purified by His TALON gravity column (Clontech) following the manufacturer’s instructions. The purified proteins were confirmed by sodium dodecyl sulfate-polyacrylamide gel electrophoresis (SDS-PAGE). OfTPSs enzyme assays were performed as described previously (Zhou et al., 2020) using GPP and FPP as substrates. After enzyme assay incubation, products were collected by the SPME for 30 mins and subsequently determined by GC-MS.

### Subcellular localization

The CDS of *OfTPS6* and *OfTPS7* without a stop codon was used to generate a C-terminal GFP fusion construct and transformed it into *A. tumefaciens* GV3101 for preparation. *Agrobacterium* cells carrying *OfWRKY33*-YFP and the Nuclear marker mCherry were mixed and cotransformed into *N. benthamiana* leaves by transient injection. Three days after injection, the fluorescence signal was observed by a confocal laser scanning microscope (TCS SP8, Leica, Germany).

### Yeast one-hybrid analysis (Y1H)

The amplified 1000 bp fragment of the *OfTPS7* promoter was cloned into HIS vector as a bait carrier, and the *O. fragrans* flowers cDNA library prey was constructed according to the CloneMiner II cDNA Library Construction Kit instruction manual. The Y1H screening was performed using the Matchmaker Gold Y1H Library Screening System (Takara, Kyoto, Japan).

To determine the interaction between OfWRKY33 and *OfTPS7*, *OfDXS1*, *OfDXR2*, and *OfIDI2*, the promoters of the above pathway genes were cloned and inserted into the pHIS2.1 plasmid to serve as reporter strains. The *OfWRKY33* ORF (1684 bp) was fused in frame with the GAL4 activation domain (AD) to generate pGADT7-OfWRKY33 as prey vector. The yeast transformants were selected on SD/-Trp /-Leu/-His medium and tested on SD/-Trp/-Leu/-His medium supplied with various concentrations of 3-amino-1,2,4-triazole (3’AT) for 2-5 d at 30 ℃. Primers used in this assay are listed in Supplementary Table S9.

### Dual-luciferase transient expression reporter assay

The PLACE signal SCAN search software (https://www.dna.affrc.go.jp/PLACE/?action=newplace) was applied to analyze the motifs of the promoters. The coding sequence of *OfWRKY33* was cloned into the pGreen II 62-SK vector functioned as an effector vector. The promoter fragments of *OfTPS7* were fused into pGreen II 0800-LUC to obtain reporter vectors. Firefly luciferase (LUC) and renilla luciferase (REN) activities were measured using dual-LUC assay kits (Promega, Madison, WI, USA). Measurements were carried out according to the manufacturer’s instructions and each measurement was repeated for six biological replicates. Primers designed for the construction of transient expression vectors are listed in Supplementary Table S9.

### Electrophoretic mobility shift assay

The CDS of *OfWRKY33* was cloned and inserted into Pet21b to obtain the recombinant vector. The recombinant vector was transformed into *Escherichia coli* BL21 (DE3) and then induced with 0.1 mmol/L IPTG (Isopropyl-beta-D-thiogalactopyranoside) at 16 °C for 16 h, followed by recombinant protein purification. Biotin-labeled probes were synthesized by Tsingke biotech company (Beijing, China). The specific sequences were presented in Supplementary Table S9. His protein was used as a control, and unlabeled probes, including identical or mutated oligonu-cleotides, were used as cold competitors. EMSA was conducted using a previously reported method (Lu et al., 2020).

### Qualitative and quantitative analysis of volatile terpenoids

The volatile analysis was based on our earlier studies (Zheng et al., 2019). Briefly, HS-SPME was employed to collect the volatile compounds from *O. fragrans* flowers and *N. benthamiana* leaves. Total volatile compounds were thermally desorbed and transferred to an GC-MS apparatus (Thermo Fisher Technologies) to analyze the compounds which were subsequently identified by comparing mass spectra with the NIST2017 mass spectra library as well as standard samples. Quantitative analysis using methyl nonanoate as internal standard. All the volatile terpenoids tests were repeated for 3 biological replicates.

### Accession number

Sequence data from this article can be found in the GenBank data libraries under accession numbers: *OfTPS6* (PQ035161), *OfTPS7* (PQ035163), *OfWRKY33* (PP598872).

Raw sequencing reads of all *Osmanthus fragrans* accessions reported in this study have been deposited into the public database of the National Center of Biotechnology Information (NCBI) BioProject under the accession number PRJNA1141249.

### Statistical analysis

The data were analyzed using GraphPad 8.0. Student’s t-test and 1-way ANOVA were conducted to determine significant differences between the 2 data sets at *P < 0.05* and *P < 0.01*. The data were expressed as mean ± SD of at least 3 biological replicates.

## Supplementary Data

The following materials are available in the online version of this article.

**Supplementary Figure 1.** Chromosomal localization of OfTPSs in *O. fragrans* ‘BYYG’ genome

**Supplementary Figure 2.** Collinearity and evolutionary types of *OfTPSs* in *O. fragrans* ‘BYYG’ genome.

**Supplementary Figure 3.** Conserved domains of OfTPSs proteins in *O. fragrans* ‘BYYG’.

**Supplementary Figure 4.** Heatmap of transcript levels of MEP and MVA pathways genes in four different blossoming stages of *O. fragrans* ‘BYYG’ flowers according to transcriptome profiles.

**Supplementary Figure 5.** Functional validation of *OfTPSs* in *N. benthamiana* leaves.

**Supplementary Figure 6.** Functional validation of OfTPS6 and OfTPS7 proteins *in vitro*.

**Supplementary Figure 7.** Subcellular localization of OfTPS6 and OfTPS7 proteins.

**Supplementary Figure 8.** Correlation analysis between potential transcription factors derived from the blue module of WGCNA and *OfTPS7*.

**Supplementary Figure 9.** Y1H analysis uncovering that OfWRKY33 does not directly interact with *OfDXS1*, *OfDXR2* and *OfIDI2*.

**Supplementary Table 1.** OAV value of dominant terpenoids aroma substances in *O. fragrans* flowers.

**Supplementary Table 2.** Statistics of the *O. fragrans* genome assembly and annotation.

**Supplementary Table 3.** Statistics of the *O. fragrans* transcriptome profiles for four different blossoming stages.

**Supplementary Table 4.** Basic information of *OfTPSs* in *O. fragrans* ‘BYYG’.

**Supplementary Table 5.** FPKM values of terpenoid synthesis related genes in the MEP and MVA pathways.

**Supplementary Table 6.** Transcript levels of five linalool synthase genes in 20 cultivars.

**Supplementary Table 7.** Contents of linalool and its oxides of *O. fragrans* flowers in in 20 cultivars.

**Supplementary Table 8.** Interaction factors obtained from *OfTPS7* through yeast single hybrid screening library.

**Supplementary Table 9.** Primer sequences utilized in this study.

**Supplementary Table 10.** The WRKY TFs from *A. thaliana*.

## Acknowledgements

We thank Prof. Deng Xiuxin (Huazhong Agricultural University, Wuhan, China) for providing informative guide for this study.

## Author contributions

R.Z. conceived and coordinated this project. W.X. and R.Z. designed the research. W.X. performed the experiments and analyzed the data with contributions from L.Z., X.Z. M.J. H.J., Q.Y., T.Y. W.X. wrote the original manuscript, R.Z. and C.W. reviewed and improved the manuscript. All authors read and approved the final manuscript.

## References

Bao TT, Shadrack K, Yang S, Xue XX, Li SY, Wang N, Wang Q, Wang L, Gao X, Cronk Q (2020) Functional characterization of terpene synthases accounting for the volatilized-terpene heterogeneity in *Lathyrus odoratus* cultivar flowers. Plant and Cell Physiology 61 (10): 1733–1749.

Benjamini Y, Hochberg Y (1995) Controlling the false discovery rate: a practical and powerful approach to multiple testing. Journal of the Royal Statistical Society. Series B (Methodological) 57: 289–300.

Bergman ME, Dudareva N (2024) Plant specialized metabolism: Diversity of terpene synthases and their products. Current Opinion in Plant Biology 81: 102607

Chen HY, Ji HY. Huang W, Zhang Z, Zhu K, Zhu S, Chai L, Ye J, Deng X (2024) Transcription factor CrWRKY42 coregulates chlorophyll degradation and carotenoid biosynthesis in citrus. Plant Physiology 195 (1): 728–744.

Chen F, Tholl D, Bohlmann J, Pichersky E (2011) The family of terpene synthases in plants: a mid-size family of genes for specialized metabolism that is highly diversified throughout the kingdom. Plant Journal 66 (1): 212–229.

Chuang YC, Hung CY, Hsu YC, Yeh CM, Mitsuda N, Ohme-Takagi M, Tsai WC, Chen WH, Chen HH (2018) A dual repeat cis-element determines expression of GERANYL DIPHOSPHATE SYNTHASE for monoterpene production in *Phalaenopsis Orchids*. Frontier in Plant Science 9: 765.

Conart C, Bomzan DP, Huang XQ, Bassard JE, Paramita SN, Saint-Marcoux D, Rius-Bony A, Hivert G, Anchisi A, Schaller H, Hamama L, Magnard JL, Lipko A, Swiezewska E, Jame P, Riveill G, Oyant LHS, Rohmer M, Lewinsohn E, Dudareva N, Baudino S, Caissard JC, Booachon B (2023) A cytosolic bifunctional geranyl/farnesyl diphosphate synthase provides MVA-derived GPP for geraniol biosynthesis in rose flowers. Proceedings of the National Academy of Sciences of the United States of America 120: 19.

Dudareva N, Cseke L, Blanc VM, Pichersky E (1996) Evolution of floral scent in Clarkia: novel patterns of S-linalool synthase gene expression in the *Clarkia breweri* flower. Plant Cell, 8 (7): 1137–1148.

Dong YM, Wei ZL, Zhang WY, Li JR, Han MX, Bai HT. Li H. Shi L (2024) LaMYC7, a positive regulator of linalool and caryophyllene biosynthesis, confers plant resistance to *Pseudomonas syringae*. Horticulture Research 11 (4): uhae044

Driguez P, Bougouffa S, Carty K, Alexander P, Kamel J, Muppala R, Richard S, Ming SC, Yoshinori F, Luca F (2021) LeafGo: Leaf to Genome, a quick workflow to produce high-quality de novo plant genomes using long-read sequencing technology. Genome biology 22 (1): 1–18.

Dudchenko O, Batra S, Omer AD, Nyquist SK, Hoeger M, Durand NC, Shamim MS, Machol I, Lander ES, Aiden AP, Aiden EL (2017) De novo assembly of the Aedes aegypti genome using Hi-C yields chromosome-length scaffolds. Science 356: 92–95.

Durand NC, Shamim MS, Machol I, Rao SS, Huntley MH, Lander ES, Aiden EL (2016) Juicer provides a one-click system for analyzing loop-resolution Hi-C experiments. Cell Systems 3: 95–98.

Fu JX, Hou D, Wang YG, Zhang C, Bao ZY, Zhao HB, Hu SQ (2019) Identification of floral aromatic volatile compounds in 29 cultivars from four groups of *Osmanthus fragrans* by gas chromatography-mass spectrometry. Horticulture Environment and Biotechnology 60 (5): 611–623.

Fu JY, Wang LP, Pei WZ, Yan J, He LQ, Ma B, Wang C, Zhu CY, Chen G, Shen QQ, Wang Q (2023) ZmEREB92 interacts with ZmMYC2 to activate maize terpenoid phytoalexin biosynthesis upon *Fusarium graminearum* infection through jasmonic acid/ethylene signaling. New Phytologist 237 (4): 1302–1319.

Gao F, Liu BF, Li M, Gao XY, Fang Q, Liu C, Ding H, Wang L, Gao X (2018) Identification and characterization of terpene synthase genes accounting for volatile terpene emissions in flowers of *Freesia* ×*hybrida*. Journal of Experimental Botany 69 (18): 4249–4265.

Gong Z, Luo YQ, Zhang WF, Jian W, Zhang L, Gao XL, Hu XW, Yuan YJ, Wu MB, Xu X, Zheng XZ, Wu GL, Li ZG, Li Z, Deng W (2021) A SlMYB75-centred transcriptional cascade regulates trichome formation and sesquiterpene accumulation in tomato. Journal of Experimental Botany 72 (10): 3806–3820.

Jiang H, Wang H (2023) Biosynthesis of monoterpenoid and sesquiterpenoid as natural flavors and fragrances. Biotechnology Advances 65: 108151.

Jiang LP, Chen S, Wang X, Sen L, Dong G, Song C, Liu Y (2024) An improved genome assembly of *Chrysanthemum nankingense* reveals expansion and functional diversification of terpene synthase gene family. BMC genomics 25 (1): 593.

Kumar S, Stecher G, Tamura K (2016) MEGA7: molecular evolutionary genetics analysis version 7.0 for bigger datasets. Molecular Biology and Evolution 33 (7): 1870–1874.

Lan YG, Zhang KM, Wang LN, Liang XY, Liu HX, Zhang XY, Jiang NQ, Wu M, Yan HW, Xiang Y (2023) The R2R3-MYB transcription factor OfMYB21 positively regulates linalool biosynthesis in *Osmanthus fragrans* flowers. International Journal of Biological Macromolecules 249: 126099.

Li HJ, Li YT, Yan HJ, Bao TT, Shan XT, Caissard JC, Zhang LS, Fang HY, Bai X, Zhang J, Wang ZX, Wang M, Guan Q, Cai M, Ning GG, Jia XJ, Boachon B, Baudino S, Gao X (2024) The Complexity of volatile terpene biosynthesis in roses: particular insights into β-citronellol production. Plant Physiology. kiae444.

Li JJ, Hu H, Mao J, Yu L, Stoopen G, Wang MJ, Mumm R, de Ruijter NCA, Dicke M, Jongsma MA, Wang CY (2019) Defense of pyrethrum flowers: repelling herbivores and recruiting carnivores by producing aphid alarm pheromone. New Phytologist 223 (3): 1607–1620.

Li SS, Zhang L, Sun M, Lv MW, Yang Y, Xu WZ, Wang LS (2022) Biogenesis of flavor-related linalool is diverged and genetically conserved in tree peony (*Paeonia* × *suffruticosa*). Horticulture Research 10 (2): 2662–6810.

Lu SW, Zhang Y, Zhu KJ, Yang W, Ye JL, Chai LJ, Xu Q, Deng XX The citrus transcription factor CsMADS6 modulates carotenoid metabolism by directly regulating carotenogenic genes. Plant Physiology 176 (4): 2657–2676.

Pichersky E, Raguso RA (2018) Why do plants produce so many terpenoid compounds. New Phytologist 220 (3): 692–702.

Roach MJ, Schmidt S, Borneman AR (2018) Purge Haplotigs: synteny reduction for third-gen diploid genome assemblies. BMC Bioinformatics 19: 460.

Shang JZ, Feng DD, Liu H, Niu LT, Li RH, Li YJ, Chen MX, Li A, Liu ZH, He YH, Gao X, Jian HY, Wang CQ, Tang KX, Bao MZ, Wang JH, Yang SH, Yan HJ, Ning GG (2024) Evolution of the biosynthetic pathways of terpene scent compounds in roses. Current Biology 34(15): 3550–3563.

Shang JZ, Tian JP, Cheng HH, Yan QM, Li L, Jamal A, Xu ZP, Xiang L, Saski CA, Jin SX, Zhao KG, Liu XQ, Chen LQ (2020) The chromosome-level wintersweet (*Chimonanthus praecox*) genome provides insights into floral scent biosynthesis and flowering in winter. Genome Biology 21(1): 200.

Stirling SA, Guercio AM, Patrick RM, Huang XQ, Bergman ME, Dwivedi V, Kortbeek RWJ, Liu YK, Sun F, Tao WA, Li Y, Boachon B, Shabek N, Dudareva N (2024) Volatile communication in plants relies on a KAI2-mediated signaling pathway. Science 383 (6689): 1318–1325.

Tholl D, Chen F, Petri J, Gershenzon J, Pichersky E (2005) Two sesquiterpene synthases are responsible for the complex mixture of sesquiterpenes emitted from *Arabidopsis* flowers. Plant Journal 42 (5): 757–771.

Wang L, Tan NN, Hu JY, Wang H, Duan DZ, Ma L, Xiao J, Wang XL (2017) Analysis of the main active ingredients and bioactivities of essential oil from *Osmanthus fragrans* var. thunbergii using a complex network approach. BMC Systems Biology 11: 144.

Wei CY, Li MT, Cao XM, Jin ZN, Zhang C, Xu M, Chen KS, Zhang B. (2022) Linalool synthesis related *PpTPS1* and *PpTPS3* are activated by transcription factor PpERF61 whose expression is associated with DNA methylation during peach fruit ripening. Plant Science 317: 111200.

Wei CY, Liu HR, Cao XM, Zhang ML, Li, X, Chen KS, Zhang B (2021) Synthesis of flavor-related linalool is regulated by PpbHLH1 and associated with changes in DNA methylation during peach fruit ripening. Plant Biotechnology Journal 19: 2082–2096.

Wu LP, Liu JY, Huang WS, Wang YX, Chen Q, Lu BY (2022) Exploration of *Osmanthus fragrans* Lour.’s composition, nutraceutical functions and applications. Food Chemistry 377: 131853.

Yang S, Wang N, Kimani S, Li YQ, Bao TT, Ning GG, Li LF, Liu B, Wang L, Gao X (2022) Characterization of terpene synthase variation in flowers of wild *Aquilegia* species from northeastern Asia. Horticulture Research 9: uhab020.

Yang XL, Yue YZ, Li HY, Ding WJ, Chen GW, Shi TT, Chen JH, Park MS, Chen F, Wang LG (2018) The chromosome-level quality genome provides insights into the evolution of the biosynthesis genes for aroma compounds of *Osmanthus fragrans*. Horticulture Research 5: 72.

Yang ZZ, Li YQ, Gao FZ, Jin W, Li SY, Kimani S, Yang S, Bao TT, Gao X, Wang L (2020) MYB21 interacts with MYC2 to control the expression of terpene synthase genes in flowers of *Freesia hybrida* and *Arabidopsis thaliana*. Journal of Experimental Botany 71 (14): 4140–4158.

Yang C, Marillonnet S, Tissier A (2021) The scarecrow-like transcription factor SlSCL3 regulates volatile terpene biosynthesis and glandular trichome size in tomato (*Solanum lycopersicum*). Plant Journal 107 (4): 1102–1118.

Yu Y, Guan J, Xu Y, Ren F, Zhang Z, Yan J, Fu J, Guo J, Shen Z, Zhao J, Jiang Q, Wei J, Xie H (2021) Population-scale peach genome analyses unravel selection patterns and biochemical basis underlying fruit flavor. Nature Communication 12 (1): 3604.

Yu ZM, Zhao CH, Zhang GH, Teixeira da Silva JA, Duan J (2020) Genome-Wide identification and expression profile of TPS gene family in *Dendrobium officinale* and the role of *DoTPS10* in linalool biosynthesis. International Journal of Molecular Sciences. 21 (15): 5419.

Zeng XL, Liu C, Zheng RR, Cai X, Luo J, Zou JJ, Wang CY (2016) Emission and accumulation of monoterpene and the key terpene synthase (TPS) associated with monoterpene biosynthesis in *Osmanthus fragrans* Lour. Frontiers in Plant Science 6: 1232.

Zhao YX, Wang MY, Chen YC, Gao M, Wu LW, Wang YD (2023) LcERF134 increases the production of monoterpenes by activating the terpene biosynthesis pathway in *Litsea cubeba*. International Journal of Biological Macromolecules 232: 123378.

Zheng RR, Liu C Wang YL, Luo j, Zeng XL, Ding HQ, Xiao W, Gan JP, Wang CY (2017) Expression of MEP pathway genes and non-volatile sequestration are associated with circadian rhythm of dominant terpenoids emission in *Osmanthus fragrans* Lour. flowers. Frontiers in Plant Science 8: 1869.

Zheng RR, Zhu ZY, Wang YL, Hu SY, Xi W, Xiao W, Qu XL, Zhong LL, Fu Q, Wang CY (2019) UGT85A84 catalyzes the glycosylation of aromatic monoterpenes in *Osmanthus fragrans* Lour. flowers. Frontiers in Plant Science 10:1376.

Zhou F, Pichersky E (2020) The complete functional characterisation of the terpene synthase family in tomato. New Phytologist 226 (5):1341–1360.

Zhou GL, Li Y, Pei F. Gong T, Chen TJ, Chen JJ, Yang JL, Li QH Yu SS, Zhu P (2022) Chromosome-scale genome assembly of *Rhododendron molle* provides insights into its evolution and terpenoid biosynthesis. BMC Plant Biology 22 (1): 342.

Zhou ZY, Xian JC, Wei WK, Xu C, Yang JF, Zhan RT, Ma DM (2021) Volatile metabolic profiling and functional characterization of four terpene synthases reveal terpenoid diversity in different tissues of *Chrysanthemum indicum* L. Phytochemistry 185: 112687.

Zhu K, Kong J, Zhao BX, Rong LX, Liu SQ, Lu ZH, Zhang CY, Xiao DG, Pushpanathan K, Foo JL, Wong A, Yu AQ (2021) Metabolic engineering of microbes for monoterpenoid production. Biotechnology Advances 53: 107837.

